# AadT, a new weapon in *Acinetobacter’s* fight against antibiotics

**DOI:** 10.1101/2023.01.03.522653

**Authors:** Varsha Naidu, Bhumika Shah, Claire Maher, Ian T. Paulsen, Karl A. Hassan

## Abstract

A novel multidrug efflux pump, AadT from the Drug:H^+^ antiporter 2 family, was discovered in *Acinetobacter* multidrug resistance plasmids. Here, we profiled the antimicrobial resistance potential and examined the distribution of this gene. Putative homologs of this efflux pump were encoded in many *Acinetobacter* species and other Gram-negative species, and were genetically associated with novel variants of *adeAB(C)*, which encodes a major tripartite efflux pump in *Acinetobacter*. The AadT pump conferred decreased susceptibility to at least eight diverse antimicrobials, including antibiotics erythromycin, tetracycline; biocides chlorhexidine; and dyes ethidium bromide and DAPI. These results show that AadT is a new determinant in the *Acinetobacter* resistance arsenal and may cooperate with variants of AdeAB(C).

## Introduction

The success of pathogenic *Acinetobacter* species in clinical settings is at least partly due to their drug resistance capabilities. Multidrug efflux pumps are significant contributors of resistance in *Acinetobacter*, as a single pump will frequently confer resistance to a broad range of antimicrobials. All *Acinetobacter* species encode efflux pumps that may confer resistance to antibiotics (1). Many of these pumps are encoded in the core genomes of the species, if not the genus, suggesting that the genes have been vertically inherited in these lineages since speciation (1-3). Others appear to be associated with mobile genetic elements, such as plasmids and transposons and have a scattered distribution in sub-sets of phylogenetically distant strains, suggesting horizontal transfer between strains (4, 5). For example, some *Acinetobacter* strains encode the tetracycline efflux pump, TetB, that was originally characterised in *Escherichia coli* on the transposon Tn*10* and is found in other Gram-negative species (6, 7).

We recently identified genes encoding a putative multidrug efflux pump from the Drug:H^+^ antiporter 2 (DHA2) family of the Major Facilitator Superfamily (MFS) of transport proteins in several *Acinetobacter* plasmids (8). Here we demonstrate that the novel DHA2 family efflux pump can decrease susceptibility to a range of antimicrobials, and that genes encoding homologs of the pump are widespread, but always associated with a variant of the *adeAB(C)* locus in *Acinetobacter* species. We call the novel efflux pump AadT, as the *Acinetobacter* <u>a</u>ntimicrobial <u>d</u>rug <u>t</u>ransporter (9).

## Methods

### Bacterial growth conditions, Cloning, PCR and Complementation

Bacterial strains were grown at 37 °C with shaking (200 rpm) in Mueller Hinton (MH) broth or LB broth or on LB plates supplemented with 100 mg/L ampicillin, and IPTG where necessary (see below). The coding region of *aadT* together with an RGSH6 epitope tag at its 3’ end was synthesised and cloned into the pTTQ18 plasmid (10), followed by next-generation sequence verification by Twist Bioscience. The recombinant plasmid carring the *amvA* gene (pTTQ18_*amvA*_) was obtained from our previous work (9). The empty plasmid (pTTQ18) and recombinant plasmids (pTTQ18_*aadt*_ and pTTQ18_*amvA*_) were transformed into chemically competent *E. coli* BL21 cells. Transformants were selected on LB plates containing 100 mg/L ampicillin and colony PCR was used to confirm positive insertion of plasmids.

### Antimicrobial susceptibility testing

Antimicrobial susceptibility assays were conducted in cation adjusted MH broth supplemented with 0.05 mM IPTG using the broth microdilution method as described (11). Assays for each antimicrobial were performed with three biological replicates and four technical replicates, resulting in twelve replicates in total. Cell growth was measured spectrophotometrically as OD_600_ endpoint readings using the PHERAstar FSX (BMG LABTECH, Germany).

### Whole Cell transport assay

*E. coli* BL21 cells carrying either pTTQ18 (empty vector), pTTQ18_*aadt*_, or pTTQ18_*amvA*_ were grown in biological replicates of three in LB broth supplemented with 100 mg/L ampicillin to OD_600_ = 0.6, followed by induction of gene expression with IPTG (0.2 mM) for 1h. The cells were washed three times in transport assay buffer (20 mM HEPES-NaOH (pH 7), 145 mM NaCl, 5 mM KCl) and resuspended in the same buffer to an OD_600_ = 1. Ethidium bromide (10 µM) and CCCP (10 µM) were added, and after 30 minutes incubation at 37 °C, the cells were washed three times in transport assay buffer and resuspended in the same buffer. Transport reaction was initiated at 37 °C by adding 1 % glucose and fluorescence was measured over time using the PTI QuantaMaster 8000 (HORIBA Scientific) with an excitation and emission wavelengths of 530 nm and 610 nm, respectively.

### Distribution of AadT and its phylogenetic relationship to other DHA2 family proteins

Variants of *aadT* were identified in NCBI databases using NCBI blast searches (see detailed descriptions in the text). Representative *aadT* loci and their surrounding genes were aligned and visualised using clicker v0.0.25 (12). The amino acid sequences of representative AadT homologs from different lineages were aligned to the sequences of other representative DHA2 family proteins using MUSCLE (13), and phylogeny inferred using MrBayes v3.2.6 (14). The pairwise sequence identity and similarity of the representative proteins was determined using MatGat (15). The same approaches were used to infer the phylogeny and sequence similarities of representative homologs of adeB.

## Results and discussion

### AadT homologs are distributed widely across the *Acinetobacter* genus

Genes encoding AadT were previously identified on plasmids carried by *A. baumannii, A. bereziniae, A. defluvii, A. johnsonii, A. nosocomialis, A. seifertii, A. ursingii* and *Acinetobacter* strains with no species designation (8). These genes were nearly identical, as only one nucleotide difference was present between sub-sets of genes, producing proteins with either a tyrosine or histidine at the predicted cytoplasmic boundary of TM helix 13. The genes were each located downstream of a gene cluster comprised of variants of *adeABC* and *adeRS* (Figure 1A), which encode a tripartite RND efflux system, AdeABC, and its two-component transcriptional regulator, AdeRS (16). The 3’ ends of *adeC* in the *aadT* linked clusters were formed by an inverted repeat sequence that was also found adjacent to a nearby IS*18* (Figure 1A) (8). The locations of the repeat sequences suggested that the intervening sequence, comprising the IS*18* transposase gene, *adeRS* and *adeABC*, may be mobilised by the IS*18* transposase. These loci also carried a partial *adeS* gene (3’ end) downstream of the *aadT* homologs, that could be related to an alternative organisation of these genes in an ancestral locus (Figure 1A).

**Figure 1.**
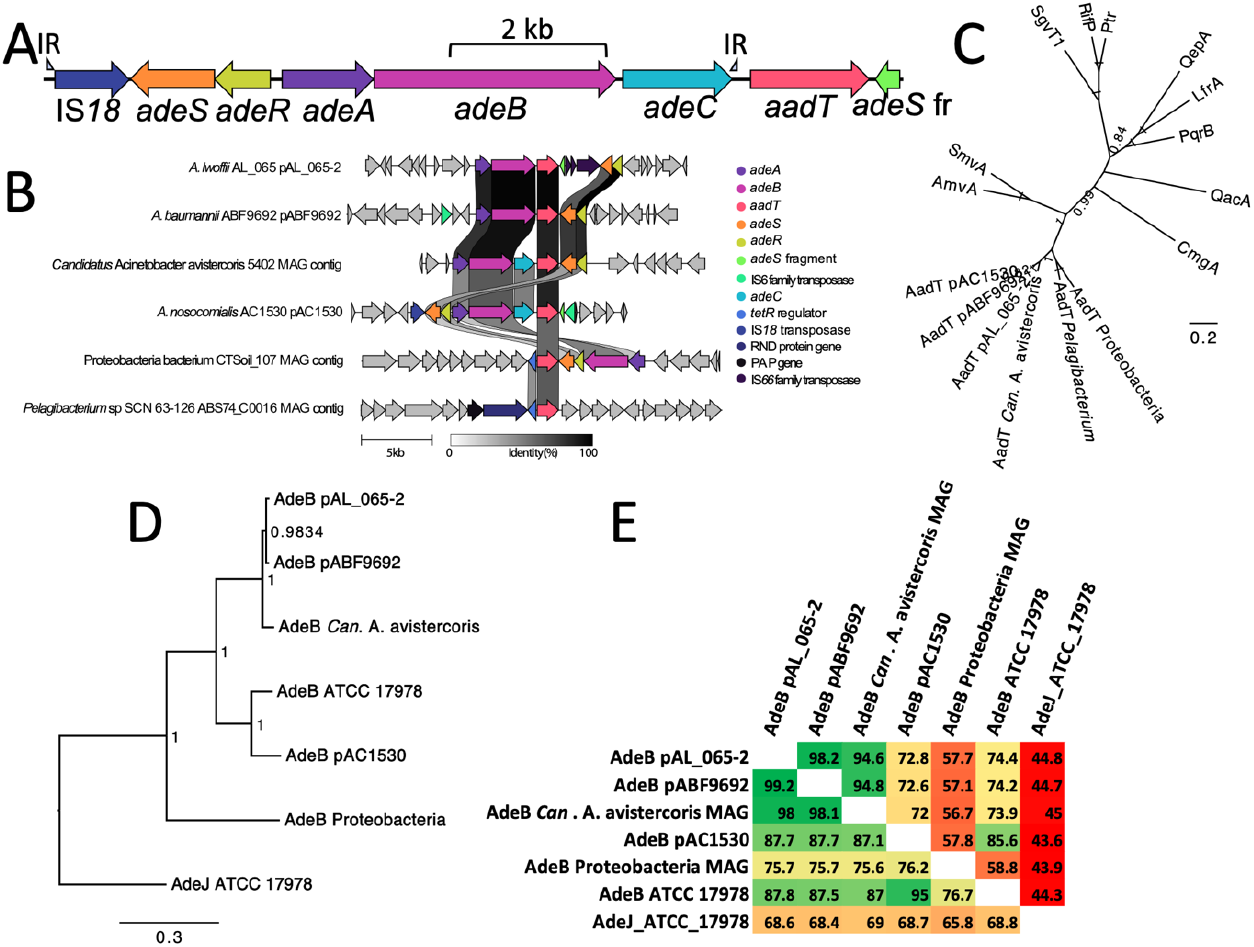
(A) Organisation of the local genetic environment of the *aadT* homolog in *A. nosocomialis* AC1530 plasmid pAC1530 and related plasmids, including the locations of surrounding genes and inverted repeat (IR) sequences that flank the IS*18* transposase gene and *adeC*. (B) Alignment of loci carrying an *aadT* homolog in selected bacterial isolates. Genes highlighted by colour and labels are found in more than one of the loci shown, or are features noted in the text. This figure was made using clinker v0.0.25 (12). (C) The amino acid sequences of AadT homologs from the lineages represented in (B) were aligned to the sequences of other representative DHA2 family proteins using MUSCLE (13), and phylogeny inferred using MrBayes v3.2.6 (14). The node labels show posterior probability values. (D) The putative AdeB homologs depicted in panel (B), and the AdeJ sequence from the *A. baumannii* ATCC 17978 chromosome were aligned using MUSCLE (13) and the phylogeny of these sequences was inferred using MrBayes v3.2.6 (14) and is shown in the tree. The node labels show posterior probability values. (E) Pairwise amino acid sequence similarity (bottom left) and identity (top right) scores for the proteins used in the phylogenetic analysis (D). These values were determined using MatGat version 2.0 (15) with a BOLSUM 50 scoring matrix.

We sought to find additional homologs of *aadT* and determine whether they are also associated with IS*18*/*adeRS*/*adeABC*. We initially queried the NCBI nr/nt database using tblastn with the AadT amino acid sequence encoded in the *A. nosocomialis* AC1530 plasmid pAC1530 and found close to 300 high scoring hits from strains designated in 23 *Acinetobacter* species and *Acinetobacter* strains with no species designation. Surveys of the local genetic environments of the high scoring hits showed one of two general arrangements. Either *aadT* was located downstream of the IS*18*/*adeRS*/*adeABC* cluster, as in pAC1530, or *aadT* was located between homologs of *adeAB* and *adeRS* gene pairs and *adeC* and the IS*18* transposase gene were absent, as in *A. baumannii* ABF9692 plasmid pABF9692 (Figure 1B). It is not unusual for *adeC* to be absent from *adeAB* loci in *Acinetobacter* genomes, and AdeAB can function with AdeK as an alternative outer membrane protein (8, 17). Insertion elements were also seen in some of the loci, such as that in *A. lwoffii* strain AL_065 pAL_065-2, where insertion sequence elements appear to have disrupted *adeS* (Figure 1B).

To identify additional homologs of AadT, we queried the NCBI experimental clustered protein database using the amino acid sequence of AadT encoded by pAC1530. This search identified 69 clusters whose representative members had above 65 % amino acid sequence identity with the query AadT sequence across at least 95 % of its length. Proteins in the top scoring cluster were encoded by *Acinetobacter* species. The local genetic environments of these homologs resembled those of *aadT* in pAC1530 or pABF9692. Other high scoring clusters included *Acinetobacter* proteins encoded by *aadT* homologs in an alternative genetic environment. For example, the genetic environment of *aadT* homologs in the *Candidatus* Acinetobacter avistercoris isolate 5402 metagenome assembled genome (MAG) and an *A. bereziniae* GD03393 plasmid resembled that in pABF9692 but included *adeC* (Figures 1B and S1). Of note, the *A. bereziniae* GD03393 plasmid harboured a second *aadT* homolog similar to pAC1530 (Figure S1).

The proteins in lower scoring clusters were encoded by species outside the *Acinetobacter* genus. The genes encoding these proteins were frequently adjacent to divergently transcribed *tetR* family regulator genes that may control expression of the *aadT* homologs. Some non-*Acinetobacter aadT* homologs were adjacent to RND efflux pump genes. For example, the *aadT* homolog in the MAG of Proteobacteria bacterium isolate CTSoil_107 (likely *Pleomorphomonas* based on GyrB and RpoD sequence similarity) was adjacent to putative distant homologs of *adeRS*/*adeAB* (Figure 1), and the *aadT* homolog from *Pelagibacterium* sp. SCN 63-126 ABS74_C0016 was encoded adjacently to a novel RND efflux system (Figure 1B). These results demonstrate a tendency for genes encoding AadT to be co-localised with those encoding RND efflux systems, specifically variants of AdeAB(C) in *Acinetobacter* species. AadT and AdeAB(C) may cooperate in the efflux of substrates from the cytoplasm to outside the cell, with AadT moving substrates from the cytoplasm to the periplasm and AdeAB(C) moving substrates from the periplasm out of the cell (18).

A phylogenetic analysis was performed to examine the relationships of AadT homologs with other members of the DHA2 family. This analysis showed that AadT homologs are most closely related to the AmvA (also known as AedF) (9, 19) and SmvA (20) efflux pumps from *Acinetobacter* and *Salmonella*, respectively (Figure 1C). Pairwise amino acid sequence similarity scores confirmed the relatedness of AadT homologs and AmvA/SmvA (Figure S2).

Phylogenetic and pairwise amino acid sequence similarity studies were conducted to examine the relationships of AdeB homologs encoded adjacently to *aadT* variants (Figure 1B). The results (Figure 1D and 1E) showed that the AdeB pump encoded in pAC1530 is most similar to the chromosomally encoded AdeB pump from *A. baumannii* ATCC 17978 (95 % amino acid sequence similarity; Figure 1E). The AdeB pumps encoded in pAL_065-1, pABF9692 and the MAG of *Candidatus Acinetobacter avistercoris* 5402 showed greater than 98 % amino acid sequence similarity to each other, but were less than 88 % similar to the chromosomal AdeB or AdeB encoded on pAC1530 (Figures 1D and 1E). The putative AdeB homolog encoded by the Proteobacteria CTSoli_107 isolate showed low similarity to all other AdeB homologs (Figure 1E). These results indicate that at least two distict putative *adeAB(C)* loci have been acquired on different *Acinetobacter* plasmids and formed an association with *aadT*. As above, the *A. bereziniae* GD03393 plasmid carries both of these loci (Figure S1).

### AadT confers resistance to multiple antibiotics, biocides and dyes

Based on the proximity of *aadT* genes to homologs of *adeAB(C)* in *Acinetobacter* species and the similarity of AadT variants to drug transporters in the DHA2 family, it seemed likely that AadT pumps would mediate resistance to one or more antimicrobials. We profiled the antimicrobial resistance potential of the AadT pump encoded in pAC1530 to eight antimicrobials and compared resistance levels to those conferred by the multidrug efflux pump AmvA in the same expression system (9, 19). The *aadT* homolog from pAC1530 was synthesised and cloned into the pTTQ18 expression plasmid (Amp^r^) behind the *tac* promoter (10) with a C-terminal RGSH6 tag and sequence verified (Twist Bioscience). The *amvA* gene from *A. baumannii* ATCC 17978 (also known as AedF) was cloned previously (9). AadT reduced susceptibility to all eight antimicrobials tested when expressed in *E. coli* BL21 cells and promoted higher levels of tolerance to tetracycline, DAPI, ethanol and ethidium bromide than AmvA (Figure 2A). These results suggest that AadT can recognise a broad spectrum of antimicrobials similar to AmvA and is a new multidrug transport protein in *Acinetobacter*.

**Figure 2.**
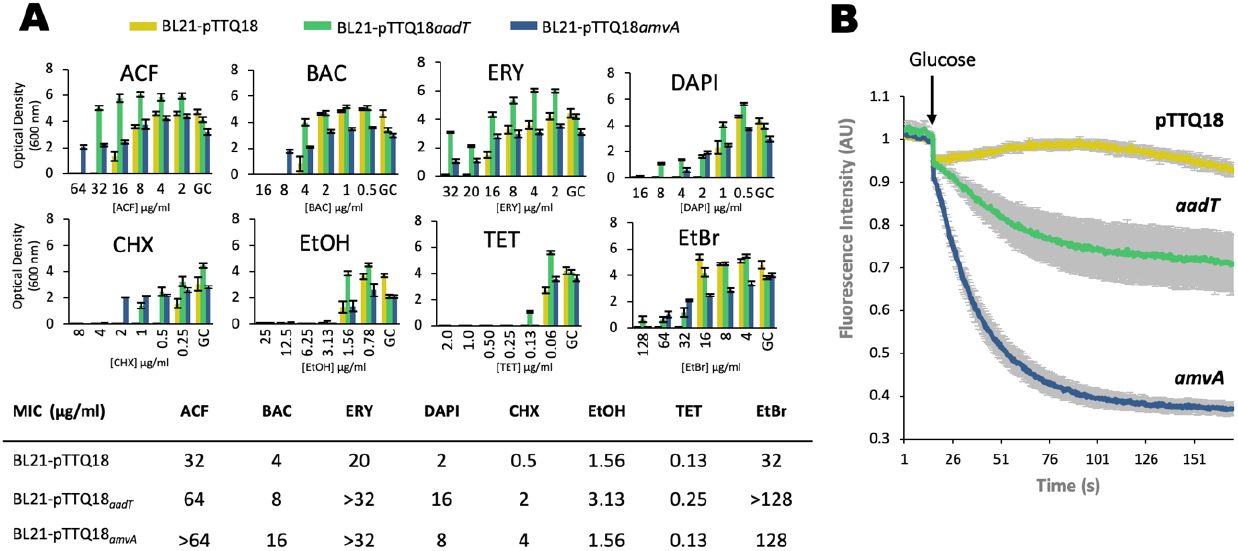
(A) *E. coli* BL21 cells carrying empty pTTQ18-*aadT* or pTTQ18-*amvA* were examined in minimum inhibitory concentration (MIC) assays for resistance to a broad range of antimicrobials, including the antibiotics tetracycline (TET) and erythromycin (ERY); the biocides benzalkonium chloride (BAC), chlorhexidine (CHX), and ethanol (EtOH); and the antimicrobial dyes 4’,6-diamidino-2-phenylindole (DAPI), acriflavine (ACR) and ethidium bromide (EtBr). The OD_600_ measurements were taken as endpoint readings after 24h growth. Resistance assays were performed as previously described (9). (B) Ethidium bromide transport assays in *E. coli* BL21 cells carrying either pTTQ18, pTTQ18-*amvA*, or pTTQ18-*aadT*. Cells were loaded with 10 µM ethidium, transport was initiated by adding glucose and efflux monitored fluorescently (Ex:450nm; Em:509nm) in a PTI QuantaMaster 8000 (HORIBA Scientific) as described previously (9). The total area under the curve (AUC) was calculcated after transport initiation by glucose for each strain BL21 pTTQ18 (150); pTTQ18-*amvA* (72); and pTTQ18-*aadT* (120). The grey shaded area represents the standard error of three biological replicates per strain.

### AadT functions via an active efflux mechanism

The ability of AadT to confer tolerance to the nucleic acid intercalating dye ethidium bromide allowed us to test efflux using real-time fluorometric transport assays. *E. coli* BL21 cells expressing AadT, AmvA or no additional protein (negative control) were loaded with 10 µM ethidium bromide in the absence of energy. The cells were washed, and re-energised and ethidium efflux monitored by fluorescence changes. Both AmvA and AadT promoted the active export of efflux ethidium bromide, as seen in fluorescence decrease over time relative to control cells (Figure 2B). This result confirms efflux as the mechanism of tolerance mediated by AadT and that AadT is a new multidrug efflux pump.

## Conclusions

The genes encoding AadT are part of the accessory genomes of more than 20 *Acinetobacter* species. This is in contrast to AmvA, its closest known relative in *Acinetobacter*, which is encoded in the core genome of *A. baumannii* and other *Acinetobacter* species. AmvA expression in *Acinetobacter* is not readily responsive to antimicrobial treatment, and recent results from our groups showed that it may have a physiological function in polyamine efflux (21). The organisation of *aadT* genes in *Acinetobacter*, downstream of *adeAB(C)* variants and in proximity to *adeRS* homologs (Figures 1A and 1B), suggests that *aadT* genes may be controlled by AdeRS and co-expressed with *adeAB(C)*. As such AadT homologs may play a more significant role in antimicrobial resistance than AmvA in the isolates that carry them. That *aadT* has formed a genetic assocication with several novel RND efflux pump genes, mostly homologs of *adeAB(C)*, suggests a selective advantage to this arrangement, which as mentioned above, may relate to cooperative transport activities of these pumps (18, 22).

The discovery of genes encoding novel variants of AdeAB(C) and AdeRS in *Acinetobacter* plasmids is significant. These genes were found in hundreds of *Acinetobacter* sequences. The variant AdeAB(C) pumps may have different substrate preferences to the chromosomally encoded pumps. Details of these substrate preferences will be of significant interest in determining their importance to antibiotic resistance in pathogenic *Acinetobacter*. The role of the plasmid encoded AdeRS system is also of significant interest. This system seems likely to control expression of the plasmid encoded *adeAB(C)* genes and as mentioned above may control *aadT* gene expression. It is also possible that the plasmid encoded AdeRS systems could promote expression of chromosomal *adeAB(C)* homologs, which are tightly regulated in many strains.

**Figure S1.**
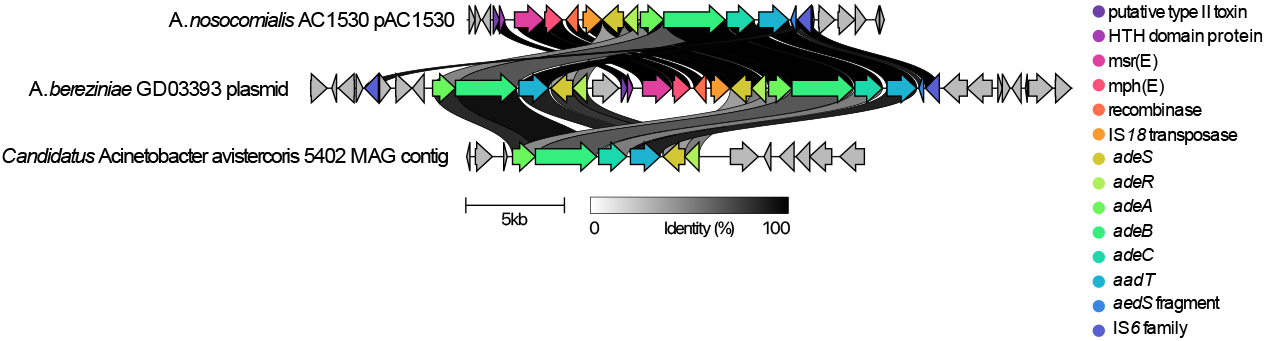
Local genetic organisation of the *aadT* loci in *A. bereziniae* GD03393 plasmid. One gene cluster shows high sequence similarity to the *aadT* locus in *A. nosocomialis* AC1530 pAC1530 and an adjacent cluster shows high similarity to the *aadT* locus in the *Candidatus Acinetobacter avistercoris* 5402 metagenome assembled genome. This figure was made using clinker v0.0.25 (12).

**Figure S2.**
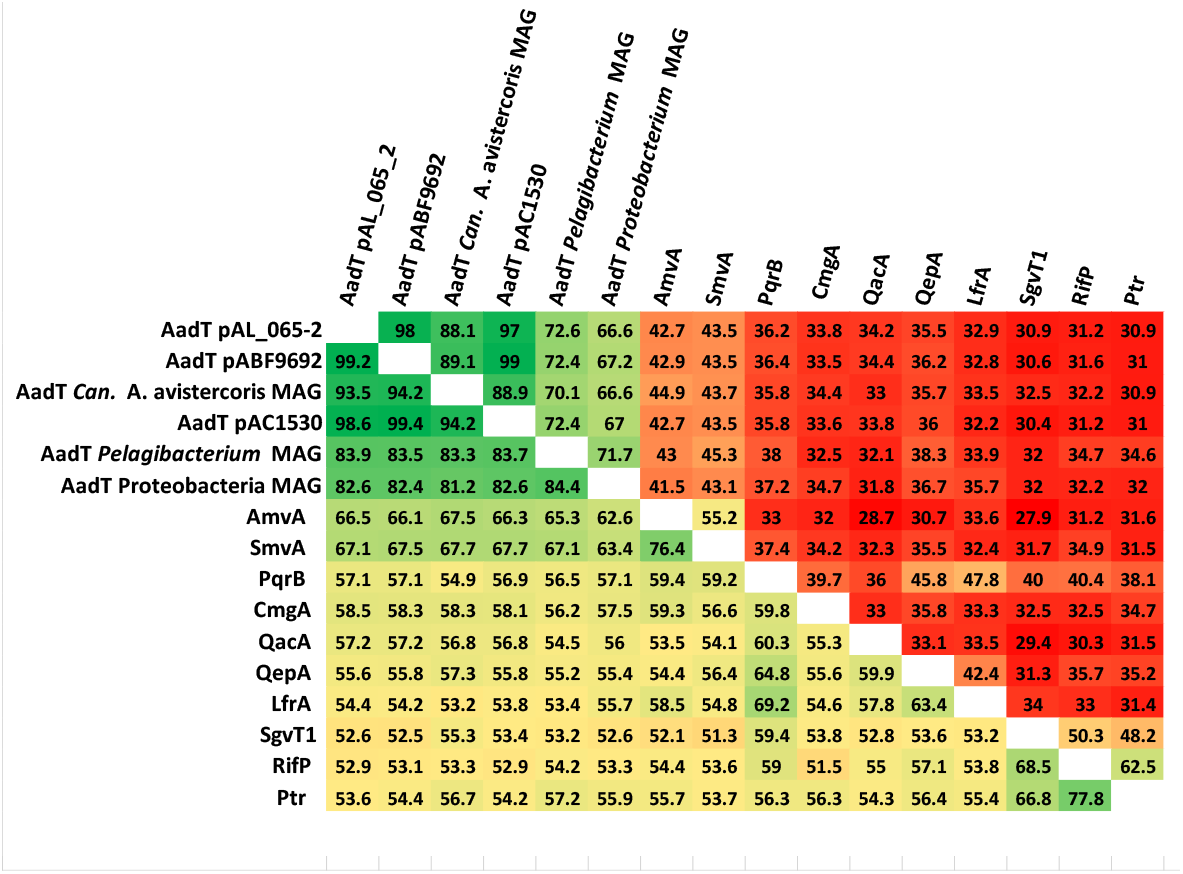
Pairwise amino acid sequence similarity (bottom left) and identity (top right) scores for members of the DHA2 family of efflux proteins. These values were determined using MatGat version 2.0 (15) with a BOLSUM 50 scoring matrix.

## Ethical Approval

N/A

